# Bioconcentration of glyphosate in wetland biofilms

**DOI:** 10.1101/2020.08.03.234856

**Authors:** Laura Beecraft, Rebecca Rooney

**Affiliations:** Department of Biology, University of Waterloo, 200 University Ave. W., Waterloo ON N2L 3G1

**Keywords:** aminomethyl phosphonic acid (AMPA), periphyton, retention, marsh, bioaccumulation, herbicide

## Abstract

Wetland biofilms were exposed to the herbicide glyphosate via *in situ* field exposures and controlled microcosm experiments to measure bioconcentration and metabolism of glyphosate by biofilm organisms. Glyphosate concentrations in biofilms were orders of magnitude higher than the surrounding water, bioconcentration factors averaged 835 and 199 in field- and lab-exposed biofilms, respectively. Glyphosate in water where it had been detected in biofilms at field-exposed sites ranged from below detection (<0.001 ppm) up to 0.13 ppm. Glyphosate bioconcentration in biofilms was inversely proportional to levels in the surrounding water, and the retention kinetics were similar to both adsorption and enzymatic models. Microorganisms present in both the water and biofilms metabolized glyphosate to its primary breakdown product aminomethyl phosphonic acid (AMPA), with increased rates of breakdown in and around the biofilms. Photosynthetic efficiency of the algae within the biofilms was not affected by 24 h glyphosate controlled exposures. Our results demonstrate the role of biofilms in improving wetland water quality by removing contaminants like glyphosate, but also as a potential exposure route to higher trophic levels via consumption. Due to bioconcentration of pesticides, exposure risk to organisms consuming or living in biofilms may be much higher than indicated by concentrations in ambient water samples.

## 1 Introduction

Since its introduction in 1974, use of the systemic, broad-spectrum herbicide glyphosate [N-(phosphonomethyl)glycine] has expanded dramatically in agriculture, silviculture, and the management of invasive plants (e.g. Annett et al., 2014; Breckels and Kilgour, 2018). Over 8.6 billion kg of the active ingredient has been applied worldwide, making glyphosate the most heavily used herbicide globally (Benbrook, 2016). Its enthusiastic adoption is attributed in part to the advent of transgenic, glyphosate resistant crops in the mid-1990s and the establishment of an inexpensive generic supply following Monsanto’s US patent expiry (Duke and Powles, 2008). Growing glyphosate use is also attributed to the development of glyphosate resistant weeds and its increasing application as a crop desiccant (Myers et al., 2016). As a result of its widespread use, glyphosate has become a ubiquitous contaminant in aquatic ecosystems (Battaglin et al., 2014; Carles et al., 2019; Lupi et al., 2019; Majewski et al., 2014; Medalie et al., 2020; Montiel-León et al., 2019).

Glyphosate works by inhibiting the enzyme 5-enolpyruvylshikimate-3-phosphate synthase (EPSPS), which blocks the shikimic acid pathway for aromatic amino acid synthesis in plants and susceptible microorganisms, including some bacteria and microalgae (Amrhein et al., 1980; Solomon and Thompson, 2003). Because the shikimic acid pathway is absent in animals (Starcevic et al., 2008), glyphosate is considered a low toxicity risk to non-target biota [e.g. 15,16]. More, glyphosate’s physicochemical properties yield a low environmental risk profile ((WHO), 2005; Giesy et al., 2000). Glyphosate is highly soluble in water (water solubility = 10,000 – 15,700□mg·L^-1^ at 25□°C; (Annett et al., 2014)), has a low octanol-water partition coefficient (log K_ow_ = −3.2; (Henderson et al., 2010)), adsorbs strongly to soil and sediment (soil adsorption coefficient = 24,000□L·kg^-1^; (Annett et al., 2014)), and can be rapidly broken down by microbial degradation (Silva et al., 2018; Solomon and Thompson, 2003). These factors contribute to a relatively short but variable half-life in water (1-91 days) (Hébert et al., 2019), and the expectation that glyphosate is rapidly dissipated from aquatic environments, with low likelihood of bioaccumulation, and minimal risk to aquatic biota (Breckels and Kilgour, 2018; Siemering et al., 2008; Solomon and Thompson, 2003).

Paradoxically, despite consistent findings of low toxicity to animals from ecotoxicology studies (Annett et al., 2014; Breckels and Kilgour, 2018; Giesy et al., 2000; Solomon and Thompson, 2003), some studies suggest that even low glyphosate concentrations may cause isruption of endocrine systems, hepatorenal damage, birth defects, teratogenic effects and alterations of the microbiome in mammals and insects (reviewed in Myers et al., 2016). Glyphosate has been shown to alter algal and bacterial abundance (Berman et al., 2020; Pizarro et al., 2016) and composition (Pérez et al., 2017; Smedbol et al., 2018) in both plankton and biofilm communities (Janßen et al., 2019; Kish, 2006; Vera et al., 2010), and it is now being recognized as a possible agent of eutrophication (Hébert et al., 2019). This is because the microbial breakdown of glyphosate releases bioavailable phosphorus (e.g. Carles et al., 2019), favoring microbes that can degrade glyphosate under low phosphorus conditions (Berman et al., 2020).

What can explain this paradox? Ecotoxicological studies of glyphosate face a variety of limitations (reviewed in Annett et al., 2014)(Annett et al., 2014). Notably, most ecotoxicology research examining the effects of glyphosate on aquatic organisms focuses on direct exposure through immersion in glyphosate contaminated water, but there may be other ecologically significant exposure pathways. For example, glyphosate adsorbs to sediment and can be taken up by both bacterial and algal cells (Forlani et al., 2008; Sviridov et al., 2015; S. Wang et al., 2016), including active and passive uptake pathways for biofilms (Battin et al., 2016). Biofilms are complex communities including bacteria, archaea, algae, viruses, fungi and protists living at the interface of substrates and the surrounding water (Battin et al., 2016; Besemer, 2015; Cui et al., 2017). The sediment and fine particles that collect in biofilms, the complex and often polyanionic nature of their exopolysaccharides, and the high-water content (Chaumet et al., 2019; Sutherland, 2001) may facilitate glyphosate bioconcentration in biofilms despite its low octanol-water partition coefficient and high water solubility. Recently, Fernandes et al. (2019) demonstrated that river biofilms in Brazil are capable of taking up glyphosate, and Klátyik et al. (2017) and Carles et al. (2019) confirmed through microcosm studies that river biofilms are capable of breaking it down. Rooney et al. (2020) observed that wetland biofilms can bioconcentrate a diverse array of agrochemicals at concentrations much higher than the surrounding ambient water. However, glyphosate was not among the pesticides examined in that study. If biofilms are bioconcentrating glyphosate from the ambient environment, biofilm grazers like invertebrates, snails, tadpoles and fish larvae might be exposed to higher concentrations of glyphosate than anticipated from water quality monitoring.

To establish whether wetland biofilms were bioconcentrating glyphosate, we employed a combination of field and controlled dose-response laboratory experiments. Our first objective was to determine the relationship between exposure dose and bioconcentration of glyphosate in wetland biofilms. In particular, we aim to test the hypothesis that if glyphosate is bioconcentrating in wetland biofilms, it is through an adsorption mechanism that would fit a saturation model. Our second objective was to assess whether wetland biofilms (i.e. the microorganisms within them) were metabolizing glyphosate to yield glyoxylate and aminomethyl-phosphonic acid (AMPA) (Sviridov et al., 2015), leading to the accumulation of AMPA in biofilms or ambient water. Our third objective was to assess if short-term glyphosate exposure affected the photochemical efficiency of the algal component of wetland biofilms, based on measurements of variable chlorophyll *a* fluorescence during the exposure period, as this could indicate cellular stress leading to metabolic changes of these autotrophs over extended exposures.

## 2 Methods

### 2.1 Biofilm colonization set-up

All biofilms were colonized *in situ* on artificial substrates comprising acrylic plates measuring 44.5 x 20.2 x 0.6 cm. These artificial substrates were installed as arrays, each consisting of 4 or 5 plates suspended vertically in the water column from marine buoys to hang ca. 10 cm below the surface of the water (Supplementary Materials Figure S1). Arrays were installed in areas of open water within the study marsh (Figure 1) at sites with 50-100 cm of standing water. These sites were selected to reduce shading from emergent or overhanging terrestrial vegetation and to avoid disturbance from boat traffic.

**Figure 1.**
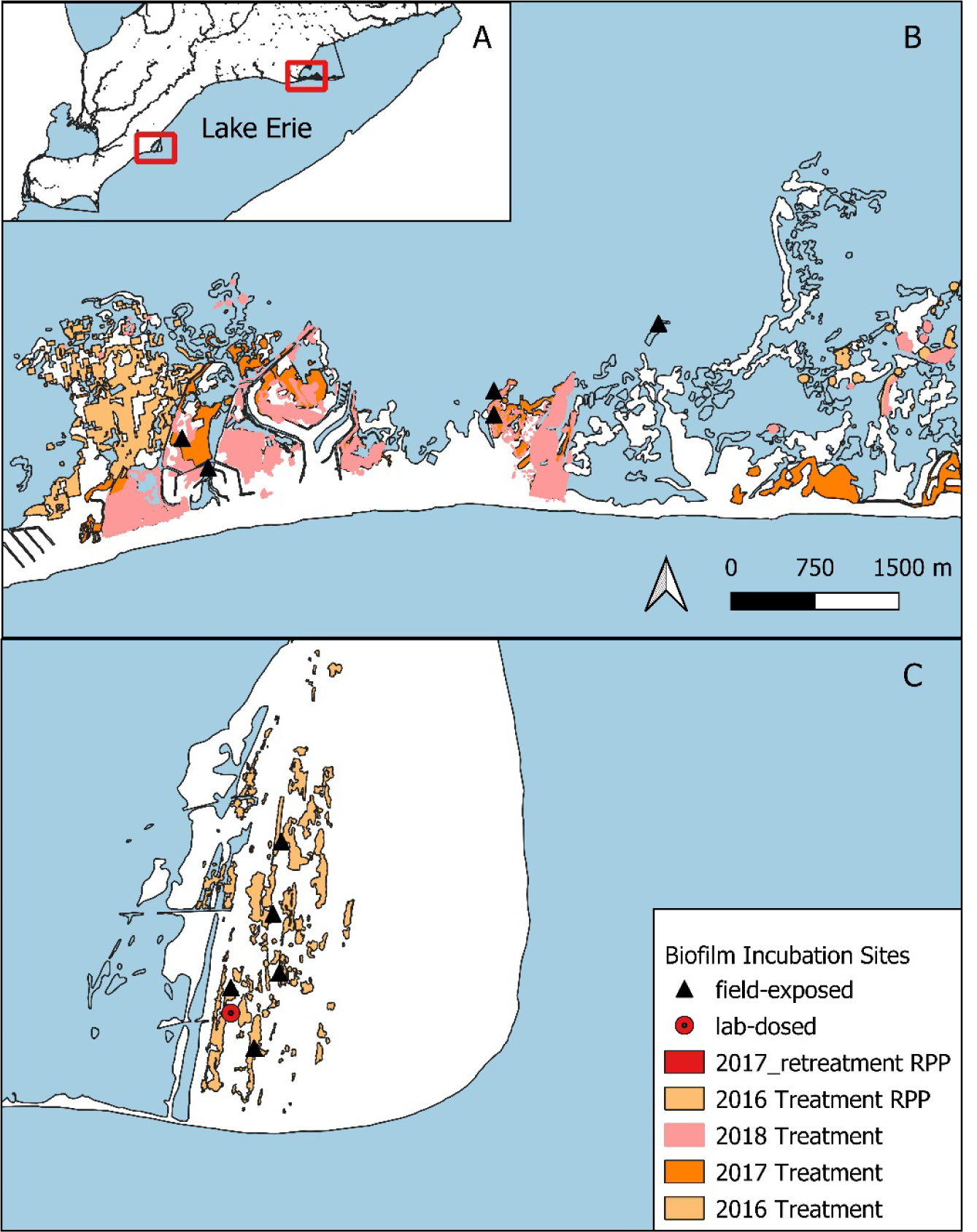
Installation sites of biofilm sampling arrays in A) two Lake Erie coastal marshes: B) Long Point Provincial Park (LPP) and C) Rondeau Provincial Park (RPP), Ontario Canada. At field-exposed sites (black triangles), biofilms colonized on artificial substrates were exposed to glyphosate applied to areas of the wetland, indicated by colour-shaded regions corresponding to treatment areas in respective years. At the ‘lab-dosed’ site (red circle, panel C), biofilms colonized on artificial substrates were collected and transported back to the lab for controlled exposures to glyphosate in microcosms.

### 2.2 Field-exposed biofilm installation and collection

Arrays were deployed in areas of open water within coastal marsh habitat in Rondeau Provincial Park (2016) and Long Point Provincial Park (2017, 2018) (Figure 1) as part of a large-scale environmental monitoring program designed around the application of glyphosate-based herbicide (Roundup Custom® for Aquatic and Terrestrial Use Liquid Herbicide, registration #32356), containing glyphosate as an isopropylamine salt with an alcohol ethoxylate surfactant (Aquasurf®, registration #32152) to control the invasive wetland grass *Phragmites australis*. The direct application of glyphosate to *P. australis* in standing water was permitted under an Emergency Use Registration from Health Canada’s Pest Management Regulatory Authority and a Permit to Perform an Aquatic Extermination from Ontario’s Ministry of Environment Conservation and Parks.

Artificial substrates were given a minimum of four weeks for *in situ* biofilm colonization prior to the date of first collection. Plates were collected from the arrays at each site on three different dates: prior to glyphosate application, 24 h after application, and approximately 40 days after application. Plates were removed from the arrays, stored in zipper-seal bags and transported in coolers back to the lab. There, we harvested the biofilm by scraping with clean cell scrapers from the plate into a Whirlpak bag and rinsing any residual biomass with small amounts of distilled/de-ionized water. All implements (scraper and funnel) were thoroughly rinsed with deionized water between samples. Due to the high level of spatial heterogeneity of the biofilms and the biomass requirements for analysis, samples from replicate plates were composited to generate a single sample per array from each sampling date and site. Samples were stored frozen and then freeze dried prior to analysis for glyphosate and AMPA by the Agriculture and Food Lab (AFL) at the University of Guelph, using the method described in ‘Chemical analyses,’ below.

At the time of plate collection in the field, we collected a depth-integrated water sample from each site using a plexiglass tube transferred to a polyethylene sample bottle, both triple-rinsed with site water. Samples were stored on ice during transport, and then frozen until delivery to AFL for analysis of glyphosate and AMPA.

This map was created using QGIS and shape files from Government of Canada, Statistics Canada, 2016 Census, in the EPSG:3347, NAD83 / Statistics Canada Lambert Projection.

### 2.3 Laboratory-Dosed biofilms

#### 2.3.1 Biofilm collection

We installed arrays containing 15 plates for the laboratory experiments in May 2018 in an open water coastal marsh site in Rondeau Provincial Park, Ontario (Figure 1C). We selected the incubation location based on 2017 surveys, where we found no detectable levels of glyphosate residue in the water or sediment. We retrieved the plates in July and transported them to the culturing facility at the University of Waterloo in coolers, placed in high-density polyethylene (HDPE) racks such that they remained upright and immersed in 100 μm-filtered lake water.

#### 2.3.2 Laboratory set up

We maintained the field-colonized biofilms in laboratory microcosms under controlled conditions. Microcosms comprised glass aquaria (ca. 37 L volume) lined with polyethylene bags to ensure glyphosate was not lost via adsorption to the glass (personal comm. from AFL to R. Rooney). Eight aquaria were filled with 100 μm filtered lake water to a volume of 28 L. The artificial substrates were held vertically (the same orientation as *in situ* colonization) in the HDPE racks. The colonized plates were distributed randomly among 5 microcosms, such that each microcosm contained 3 plates. The remaining 3 microcosms contained filtered lake water and 3 clean, un-colonized, plates each, which we used as experimental controls to account for glyphosate loss and/or metabolism occurring in the filtered lake water itself, and possible adsorption of glyphosate to the acrylic plates. We left the microcosms for 72 h to equilibrate to laboratory growth conditions: 40 μmol photons ·m^2^·s^-1^ at water surface from cool white fluorescent lights under a 16:8 hr light: dark cycle and constant aeration from air pumps and diffuser tubes (Supplementary Materials Figure S2). Temperature and dissolved oxygen were maintained at 21 ± 1 °C and 8.7 ± 0.1 mg·L^-1^ (ca. 100%), respectively, though dissolved oxygen briefly reached 70-80% saturation in the coolers during transport to the culturing facility. Water level was maintained at 28 L with additions of filtered lake water to replaces losses from evaporation.

#### 2.3.3 Glyphosate exposure

The microcosms were exposed to different concentrations of glyphosate for 24 h in a regression design. Treatments for microcosms containing colonized plates (‘colonized microcosms’) had nominal glyphosate concentrations of 0, 0.01, 0.1, 1.0 and 10 mg glyphosat a.e. · L^-1^, respectively, and 0, 0.1 and 10 mg glyphosate a.e. · L^-1^ for the microcosms containing clean plates (‘control microcosms’). These exposure levels were chosen to encompass concentrations observed in natural surface waters by other researchers (Annett et al., 2014; Battaglin et al., 2014) and to create a gradient from which to assess glyphosate bioconcentration. To achieve the desired exposure levels, we added glyphosate from a stock solution (480 mg glyphosate a.e. · L^-1^) made from a dilution of RoundUp Custom® (original concentration of 480 g glyphosate a.e. · L^-1^).

After 24 h, we collected water samples from each microcosm to compare measured and nominal concentrations. Samples were taken in acid washed polyethylene sample bottles, rinsed in triplicate with sample water. We harvested the biofilms from the plates using scraping tools (plastic putty knives), rinsed thoroughly with de-ionized water between samples. Biofilms from the three plates in each tank were composited, transferred to Whirlpak bags and frozen. Samples were freeze-dried at −50°C. Water and biofilm samples were stored frozen until delivery to AFL for analysis of glyphosate and AMPA.

During the 24 h glyphosate exposure period, we used a pulse-amplitude modulated chlorophyll *a* (Chl *a*) fluorometer (Diving-PAM, Walz Effeltrich Germany) to measure the quantum yield of photochemistry (ΔF/F_m_ ^’^) of the photosynthetic organisms within the biofilms, to detect stress responses of Photosystem II photochemistry due to glyphosate exposure. The Diving-PAM measures background fluorescence (F) using low intensity, non-actinic, modulated red light (655 nm LED), followed by a saturating pulse of red light, which oxidizes all reaction centers and induces maximum fluorescence (F_m_’) (Hiriart-Baer et al., 2008; Walz GmbH, 1998). We measured both background and maximum fluorescence in the light-adapted state, as it was not possible to dark-adapt the biofilms on the plates while taking replicate measurements and without removal from their respective treatment microcosms. ΔF/F_m_ ^’^ is calculated as (F_m_ ^’^ – F)/ F_m_ ^’^ (Cosgrove and Borowitzka, 2010). A magnetic sample holder was attached to the fiber-optic sensor to ensure the sensor remained a constant distance from the sample during measurement. Ten replicate measures at different locations were taken on each of the three colonized plates for each treatment, starting with pre-exposure (time 0) and then 0.5, 1, 2, 3, 6 and 24 h post-dose, rinsing the sensor thoroughly between microcosms.

### 2.4 Chemical Analyses

The Agriculture and Food Laboratory (AFL) at the University of Guelph conducted the analyses of glyphosate and AMPA for all water and biofilm samples (limits of detection, Supplementary Materials Table S1). Samples were first homogenized, fortified with internal standard and then centrifuged. The supernatant was then acidified prior to liquid chromatography and mass spectrometry, and the samples quantified using a ratio of external to internal standard. Results are reported in ppm, equivalent to mg·L^-1^.

### 2.5 Statistical Analyses

Linear regression analyses were used to determine how AMPA concentration in biofilm tissues or water depends on the glyphosate concentration in that same substrate. The adsorption of glyphosate to the biofilm from the surrounding water was modelled using both adsorption (1) and enzyme (2) kinetics. We considered an adsorption and enzymatic model because both processes may be occurring in the glyphosate-biofilm interaction; glyphosate is adsorbing to the different biofilm components, but is also being taken up and metabolized by cells/organisms within the biofilm. The Freundlich adsorption isotherm (1) is an empirical relationship between solute adsorbed to a surface and solute in the surrounding liquid, which can be expressed as:

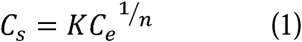

Where *C*_*s*_ is the concentration of adsorbed herbicide, *C*_*e*_ is the herbicide concentration in the surrounding water at equilibrium, *K* is the Freundlich adsorption constant, and *1/n* is a constant relating adsorption to pressure (Alister et al., 2010). The Freundlich absorption isotherm can be determined in relation to either equilibrium pressure or concentration of the absorbate; we are using the latter and assuming that equilibrium concentrations of glyphosate in the microcosms had been reached at 24 h, supported by previous studies of pesticide accumulation in biofilms (Chaumet et al., 2019; Lundqvist et al., 2012).

The Michaelis-Menten equation models enzyme kinetics by relating enzyme reaction rates to substrate concentration, and is expressed as:

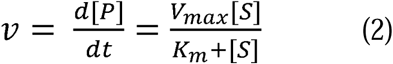

Where *v* is reaction rate, *S* is the substrate (glyphosate is water), *P* is the product (glyphosate adsorbed in biofilm), *V*_*max*_ is the maximum reaction rate achieved by the system, and *K*_*m*_ is the Michaelis constant – the substrate concentration at which the reaction rate reaches half of maximum.

Bioconcentration is the retention of a substance in an organism’s tissues relative to the surrounding environment, taken up by contact and respiration (Alexander, 1999; Arnot et al., 2006). Bioconcentration factor (BCF) was calculated on a dry-weight basis by dividing the concentration of glyphosate (or AMPA) in the biofilm (dry-weight) by that in the surrounding water, assuming a steady state had been reached after 24 h of exposure. For field sites where glyphosate and/or AMPA was detected in biofilm but not in the water, we used the limit of detection (or quantification) (Supplementary Materials Table S1), as appropriate, for the water concentration, providing a conservative estimate of the BCF. The dependence of BCF on ambient water concentrations fit a power function relationship, which we quantified using linear regression on the log-transformed values. A general linear model of the log-transformed values was used to assess if the relationship was significantly different based on application type (i.e. field-exposed vs. lab-dosed) or chemical (glyphosate vs. AMPA) (Supplementary Materials Table S2). In both cases the factors did not have a significant main or interaction effect and a single regression was used to model the relationship.

We assessed the relationship of AMPA concentration in biofilm or water to glyphosate in that same substrate by linear regression. Slope was estimated with an intercept estimate and with the intercept set to zero. We reported regression parameters for the latter for three reasons: analysis of variance (ANOVA) indicated no significant difference between the linear regression models with and without intercept estimates; models with an intercept predicted negative AMPA concentrations at low glyphosate concentrations; and an intercept of zero is the logical format from a biological/chemical perspective. A general linear model was used to confirm that there was no significant difference in the AMPA-glyphosate relationship between field-exposed and lab-dosed biofilms (Supplementary Materials Table S3) and so the regression was estimated for lab-dosed and field-exposed biofilms combined.

The effect of glyphosate exposure on the quantum yield of photochemistry, ΔF/F_m_ ^’^, was assessed by linear regression of the change in ΔF/Fm’ post-exposure, normalized to initial values (i.e. (post-exposure – pre-exposure) / pre-exposure). Statistical analyses were performed in Excel and R Statistical Software version 4.0.1 (R Core Team, 2020), including the packages tidyverse (Wickham et al., 2019), rstatix (Kassambara, 2020) and broom (Robinson and Hayes, 2020).

## 3 Results

The uptake of glyphosate from the surrounding water into the biofilm tissues followed a power function relationship, which we modelled using the Freundlich adsorption isotherm and Michaelis-Menten enzyme kinetics (Figure 2). Glyphosate and AMPA bioconcentrated in biofilm tissue by two to three orders of magnitude relative to the surrounding water, with BCF_DW_ ranging from 11 to 23,500 for glyphosate and from 4 to 3200 for AMPA (Table 1). The BCF_DW_ of glyphosate and AMPA were strongly dependent on the herbicide concentration in the ambient water, following a negative power function relationship (F_1, 20_ = 39.62, p < 0.0001) (Figure 3). This relationship was not significantly different between lab-dosed and field-exposed biofilm samples, based on a two-factor general linear model (p = 0.903, Supplementary Materials Table S2).

**Figure 2.**
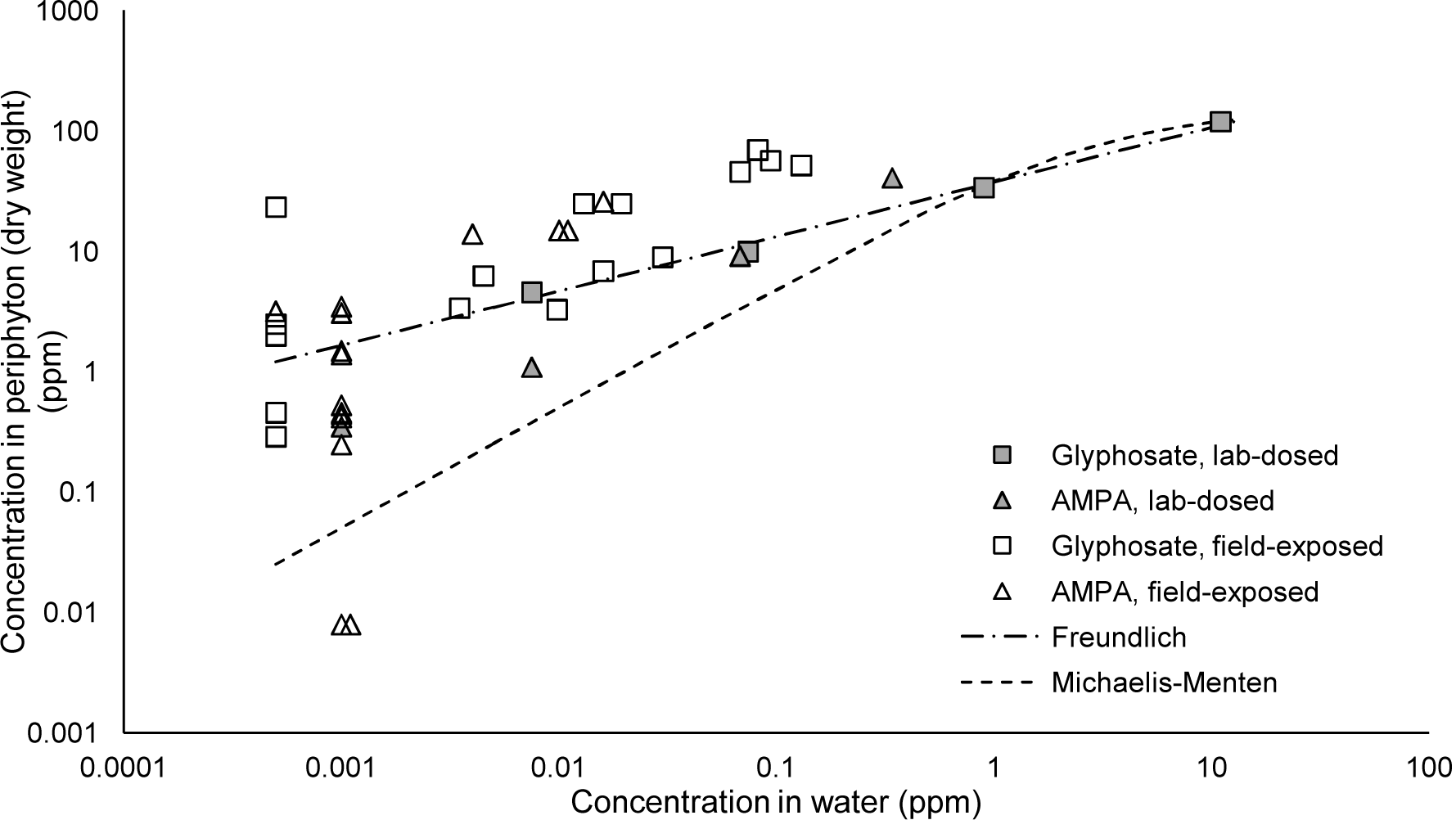
Glyphosate and AMPA retention in biofilms relative to concentrations in the surrounding water. Models of glyphosate retention kinetics in biofilm tissues were estimated from lab-dosed biofilms in microcosm experiments (grey symbols). We used the Freundlich adsorption isotherm: *C*_*s*_ = 37.497*C*_*e*_^1/2.21^ (dashed line) and Michaelis-Menten enzyme kinetics (*d[P])****/****dt*=(156.056[*S*])/(3.039+[*S*]) (dotted line).

**Table 1.** Dry-weight bioconcentration factors (BCF_DW_) of glyphosate and aminomethyl phosphonic acid (AMPA) in biofilms relative to the surrounding water from field-exposed and lab-dosed (‘lab’) biofilms. BDL and BQL indicate herbicide concentrations below methodological detection or quantification limit, respectively. In these cases, the limit of detection or quantification (LOD/LOQ) in water (Table S1) was used to conservatively estimate the BCF, however these samples were not included in calculation of the average BCFs.

| Wetland Site | Glyphosate Treatment | Water | | Periphyton | | BCF | | Average BCF (± SD) | |
| --- | --- | --- | --- | --- | --- | --- | --- | --- | --- |
|  |  | Glyphosate (ppm) | AMPA (ppm) | Glyphosate (ppm dw) | AMPA (ppm dw) | Glyphosate | AMPA | Glyphosate | AMPA |
| RPP - 2016 | field-exposed | 0.068 | 0.016 | 46.0 | 26.0 | 676 | 1625 |  |  |
| RPP - 2016 | field-exposed | 0.130 | 0.010 | 52.0 | 15.0 | 400 | 1500 |  |  |
| RPP - 2016 | field-exposed | 0.013 | BDL | 25.0 | 3.2 | 1923 | 3200 |  |  |
| RPP - 2016 | field-exposed | 0.094 | 0.011 | 57.0 | 15.0 | 606 | 1364 |  |  |
| RPP - 2016 | field-exposed | 0.082 | BQL | 70.0 | 14.0 | 854 | 1750 |  |  |
| LPP - 2017 | field-exposed | 0.019 | BDL | 25.0 | 3.5 | 1291 | 1750 |  |  |
| LPP - 2017 | field-exposed | BDL | BDL | 0.5 | BDL | 460 | 4 |  |  |
| LPP - 2017 | field-exposed | BDL | BDL | 0.3 | BDL | 290 | 4 | 835 ± 519 | 1496 ± 131 |
| LPP - 2017 | field-exposed | BDL | BDL | 23.5 | 3.1 | 23500 | 1550 | (n=11) | (n=3) |
| LPP - 2018 | field-exposed | 0.0035 | BDL | 3.4 | 0.4 | 971 | 210 |  |  |
| LPP - 2018 | field-exposed | 0.016 | BDL | 6.9 | 0.3 | 431 | 125 |  |  |
| LPP - 2018 | field-exposed | BDL | BDL | 2.5 | 0.5 | 2500 | 230 |  |  |
| LPP - 2018 | field-exposed | 0.03 | BDL | 9.0 | 1.5 | 300 | 750 |  |  |
| LPP - 2018 | field-exposed | 0.0098 | BDL | 3.3 | 0.5 | 337 | 265 |  |  |
| LPP - 2018 | field-exposed | BDL | BDL | 2.0 | 0.5 | 2000 | 225 |  |  |
| LPP - 2018 | field-exposed | 0.0045 | BDL | 6.3 | 1.4 | 1400 | 700 |  |  |
| RPP - 2018 | lab (0.01 ppm) | 0.008 | 0.001 | 4.6 | 0.4 | 613 | 175 |  |  |
| RPP - 2018 | lab (0.1 ppm) | 0.074 | 0.008 | 10.0 | 1.1 | 135 | 147 | 199 ± 281 | 134 ± 13 |
| RPP - 2018 | lab (1.0 ppm) | 0.900 | 0.068 | 34.0 | 9.2 | 38 | 135 | (n=4) | (n=3) |
| RPP - 2018 | lab (10 ppm) | 11.000 | 0.340 | 120.0 | 41.0 | 11 | 121 |  |  |

Note: Average BCF values do not include samples where water concentrations were BDL or BQL to provide a conservative estimate of average BCF.

**Figure 3.**
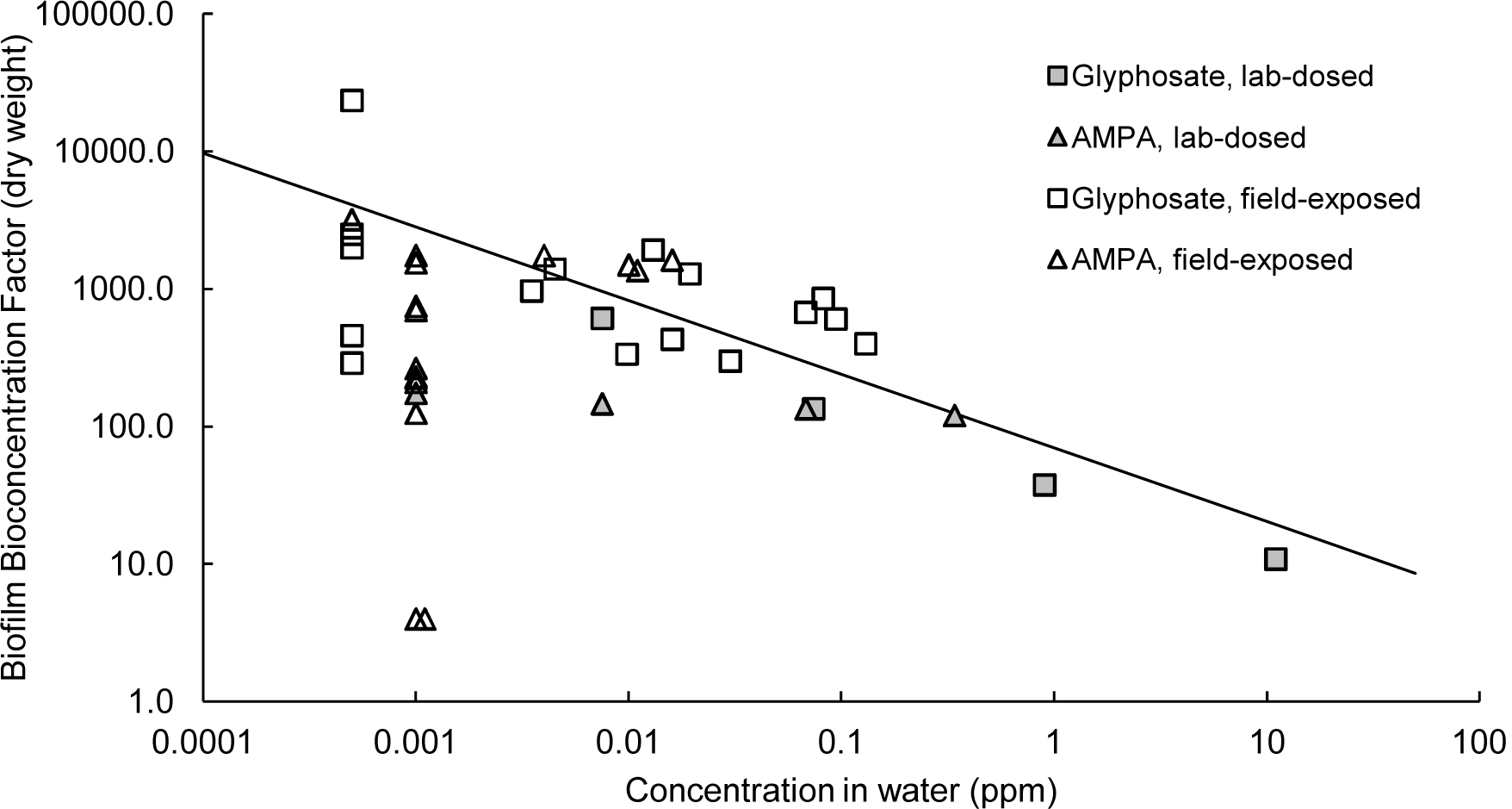
Dry-weight bioconcentration factors (BCF_DW_) of glyphosate and AMPA in biofilm tissues from microcosms (lab-dosed) and field-exposed samples vary with ambient water concentration in a power function relationship: *y* = 69.88 · *x*^-0.536^, residual standard error = 0.3564, F_1, 20_ = 39.62, p < 0.0001.

The concentration of AMPA was strongly and significantly dependent on the concentration of glyphosate in microcosm water (Figure 4a) and biofilm material (Figure 4b), with much greater regression slopes in the biofilms compared to the filtered lake water (Table 2). The rate of glyphosate metabolism to AMPA (i.e. regression slope) was not significantly different between lab-dosed and field-exposed biofilms, based on a two-factor general linear model (p = 0.705, Supplementary Materials Table S3).

**Table 2.** Linear regression analysis of the AMPA-glyphosate relationship in biofilms and microcosm water, including control microcosms containing un-colonized plates, with regression

| Medium | Linear regression results |  |  |  |  |  |
| --- | --- | --- | --- | --- | --- | --- |
|  | Slope | df | RSE | F-statistic | p-value | R<sup>2</sup> |
| Biofilm | 0.2997 | 28 | 3.4260 | 240.2 | 2.87E-15 | 0.8919 |
| Water (colonized microcosms) | 0.0312 | 4 | 0.0202 | 291 | 6.93E-05 | 0.9831 |
| Water (control microcosms) | 0.0092 | 2 | 0.0036 | 941.5 | 0.00106 | 0.9968 |

parameters including degrees of freedom (df), residual standard error (RSE) and adjusted R<sup>2</sup>.

**Figure 4.**
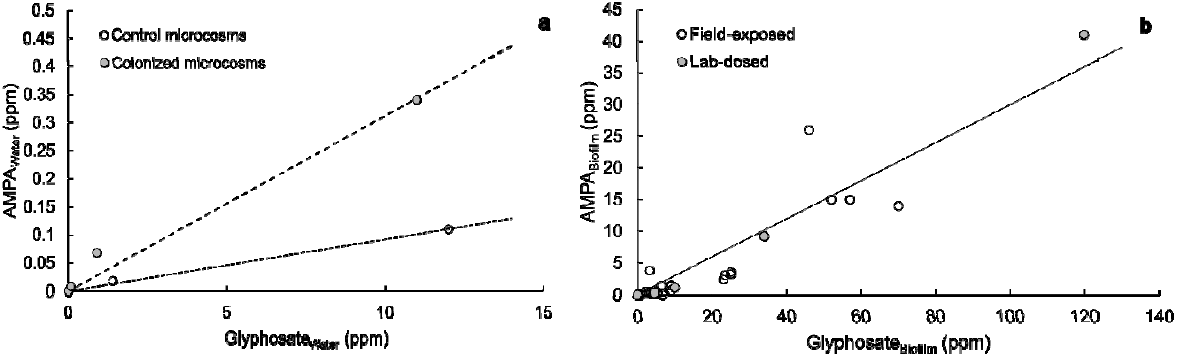
Concentration of the breakdown product, AMPA, increases linearly (Table 2) with glyphosate concentration in (**a**) water of control (open symbols) and colonized (grey symbols) microcosms after 24 h exposure, and in (**b**) biofilm tissues dosed for 24 h under laboratory conditions (‘lab-dosed’, grey symbols) and biofilms exposed *in situ* for 24 h – ca. 40 days (‘field-exposed’, open symbols).

Algal abundance and composition were heterogeneous within the biofilms, based on the variability in replicate ΔF/F_m_’ measurements taken from each plate and across plates within a given microcosm (Supplementary Materials Table S4), though pre-exposure ΔF/F_m_’ was not significantly different between microcosms based on one-way ANOVA (F_4, 166_ = 2.307, p = 0.060). There was a non-significant linear relationship between the normalized change in post-exposure ΔF/F_m_’ compared to glyphosate exposure concentration (y = −0.0068*x* + 0.0755, F_1,3_ = 2.628, p = 0.2034, adjusted r^2^ = 0.2893; Supplementary Materials Figure S3), suggesting acute (24 h) exposure at the concentrations tested causes little to no decrease in the efficiency of photochemistry (Photosystem II activity) of the algal/photosynthetic component of the biofilms.

## 4 Discussion

Biofilms are ecologically important for a number of reasons, including that they adsorb, retain and amplify solutes, accumulating substances that are otherwise highly dilute in the surrounding water (Battin et al., 2016; Sabater et al., 2002), with evidence of biofilms accumulating herbicides (Chaumet et al., 2019; Klátyik et al., 2017; Lawrence et al., 2001; Nikkila et al., 2001), insecticides (Lundqvist et al., 2012), PCBs (Wang et al., 1999), and a variety of other pesticides (Mahler et al., 2020; Rooney et al., 2020). Glyphosate is the most heavily used herbicide globally and is accumulating in our wetlands. Although its direct toxicity to fauna is well characterized as low risk (e.g. Giesy et al., 2000), there is a growing body of literature documenting the indirect effects of chronic glyphosate exposure to a wide range of aquatic organisms (Florencia Gutierrez et al., 2017; Myers et al., 2016; Pizarro et al., 2016; Vera et al., 2010). Our first objective was to determine if bioconcentration of glyphosate occurs in wetland biofilms, and test how the bioconcentration factor varies with exposure dose. Our second objective was to assess whether the glyphosate interacting with wetland biofilms is available for metabolism and to measure the extent of breakdown of glyphosate to AMPA by biofilm organisms.

We observed retention and bioconcentration of glyphosate and its breakdown product AMPA in lab-dosed and field-exposed biofilms. The herbicide had strong adsorption at low ambient concentrations, and an apparent saturation effect at higher ambient concentrations, similar to observations of diuron accumulation by Chaumet et al. (2019). This is supported by the strong fit of both Michaelis-Menten enzyme kinetics and Freundlich adsorption models to our data. Furthermore, observed BCFs more closely followed the Freundlich adsorption isotherm at low ambient water concentrations (< 1 ppm) and Michaelis-Menten kinetics at higher concentrations (> 1 ppm). Biologically, this may correspond to initial, rapid adsorption of herbicide to biofilm surfaces and the extracellular polymeric substances of the biofilm matrix, followed by slower enzymatic uptake and metabolism of the herbicide by biofilm microorganisms. The result is a BCF inversely proportional to glyphosate concentration in the surrounding water.

The BCFs of glyphosate and AMPA in field-exposed biofilms were higher than in lab-exposed biofilms, likely because the observed herbicide concentrations in the water were lower in the field compared to the laboratory-dosed microcosms. However, the relationship of BCF to ambient water concentrations did not differ significantly between field-exposed and lab-dosed biofilms, or between glyphosate and AMPA, suggesting that the microcosms captured the same mechanisms important to bioconcentration in the wetlands. Thus, although the levels of glyphosate and AMPA detected in natural surface waters are typically quite low (e.g., 0.159 μg glyphosate·L^-1^ (Glozier et al., 2012); < 0.03 μg glyphosate·L^-1^ (Annett et al., 2014; Battaglin et al., 2014), 0.1-0.3 μg glyphosate·L^-1^ (Carles et al., 2019)), the glyphosate concentration in biofilm tissues may be much higher due to bioconcentration, with BCFs greater than 800 at concentrations reported to be typical of surface waters.

Our results contradict the expectation that glyphosate will not bioaccumulate or bioconcentrate based on its chemical characteristics of high solubility, low octanol-water partition coefficient, and ability to be broken down by environmental microorganisms (Breckels and Kilgour, 2018; Solomon and Thompson, 2003). On the other hand, bioconcentration in biofilms offers an explanation for the apparent rapid dissipation of glyphosate from surface waters (Goldsborough and Brown, 1993); e.g., like us, Klátyik et al. (2017) observed the accelerated dissipation of glyphosate from river water in microcosms containing biofilms within 24 h of herbicide addition. Glyphosate bioconcentration has been observed in other organisms: leaf tissues of the aquatic macrophyte *Ludwigia peploides* from surrounding surface waters (Pérez et al., 2017), and in the oligochaete *Lumbriculus variegatus*, with the uptake/adsorption relative to water concentration fitting the Freundlich adsorption isotherm (Contardo-Jara et al., 2009). Glyphosate concentrations in surface water can spike immediately after application and runoff events, and then drop very rapidly, with concentrations much lower in water collected only a few hours later (Goldsborough and Beck, 1989; Peruzzo et al., 2008; Robichaud and Rooney, n.d.). The rapid adsorption and bioconcentration by biofilms may both contribute to this rapid removal, as well as retain the glyphosate in a given environment or location longer than was previously realized, by retaining it within the biofilm tissues rather than in the water. It would be useful to know the rate and extent of depuration of glyphosate and AMPA from biofilms back into the surrounding water column as herbicide concentrations in the water decrease, which would inform potential flushing of glyphosate from biofilms after herbicide exposure.

Glyphosate is metabolized by a variety of microorganisms in soil, water and sediment (Solomon and Thompson, 2003; S. Wang et al., 2016), including organisms that can reside in biofilms. Furthermore, wetlands have been shown to facilitate biodegradation of glyphosate to AMPA in runoff, correlated with the presence of resident wetland vegetation (Imfeld et al., 2013; Liu et al., 2019). The wetland biofilms used in the present study metabolized glyphosate from the surrounding water, consistent with observations from other studies (Carles et al., 2019; Klátyik et al., 2017). The stable structure of a biofilm allows for the formation of a functional community that is more dense and metabolically efficient compared to planktonic cells (Besemer, 2015). When comparing the dependence of AMPA concentration to glyphosate concentration in biofilm and water samples, linear regression slopes were highest for the biofilms themselves, followed by water from microcosms containing colonized plates, and lowest in water from control microcosms. This indicates that microorganisms present in both the 100 μm-filtered lake water and the biofilms are metabolizing glyphosate to AMPA, but that the biofilms are primarily responsible for glyphosate metabolism and are releasing AMPA into the surrounding water. The rate of conversion (slope) from glyphosate to AMPA was not significantly different between the lab-dosed and field-exposed biofilms, and was two orders of magnitude higher within the biofilms compared to filtered lake water. Thus, we conclude that biofilms increase the rate and extent of glyphosate metabolism in their environment, as suggested by Lawrence et al., (2001) and Klatyik et al. (2017). These results offer a possible explanation for the observations of Imfeld et al. (2013) that transport and degradation of glyphosate in stormwater wetlands are influenced by the vegetation: increased vegetation may have provided increased surface area for biofilm colonization, and retention and degradation of the biofilms facilitated the observed changes in water concentrations.

Glyphosate and its breakdown product AMPA are frequently detected in surface waters of Canadian streams and rivers (e.g. Struger, Stempvoort, and Brown 2015) usually at levels below the Canadian Water Quality Guideline for the protection of aquatic life (800 μg glyphosate·L^-1^ for chronic exposure, (CCME, 2012)). However, if concentrations in biofilms are orders of magnitude higher than surface waters, it introduces a dietary exposure route to grazing organisms via consumption (Lundqvist et al., 2012). Despite considerable research on organism sensitivity to glyphosate-based herbicides, there is also disparity in the reported results. Toxicity responses vary with species, exposure route and duration (Annett et al., 2014), ranging from: negligible (reviewed in Breckels and Kilgour, 2018)(Breckels and Kilgour, 2018); to moderate (e.g. cladoceran (Tsui and Chu, 2003); snails (Druart et al., 2017); amphibians (Carvalho et al., 2018; Druart et al., 2017); fish (Zebral et al., 2018)); to strong negative impacts (e.g., amphibians (King and Wagner, 2010; Paganelli et al., 2010; Relyea and Jones, 2009)). This apparent discrepancy over the magnitude of risk that glyphosate poses to aquatic biota remains because the effects of glyphosate-based herbicides on non-target aquatic organisms differ by dose, exposure route, timing of exposure and taxon studied (Annett et al., 2014; Reno et al., 2014; Tsui and Chu, 2003). It is further complicated because the commercially available glyphosate formulations comprise a proprietary blend of constituents to improve herbicide efficacy (Druart et al., 2017; Klátyik et al., 2017; Myers et al., 2016). Some studies have attributed the toxicity of glyphosate-based herbicide formulations to these additives, rather than the glyphosate *per se* (e.g. Reno et al., 2014; Tsui and Chu, 2003), yet these additives may be challenging to identify and differ among products, hampering synthesis of the toxicological literature on glyphosate (reviewed in Annett, Habibi, and Hontela 2014). Thus, the formulation and concentration in acid equivalents need to be considered in studies assessing risk to aquatic life.

Bioconcentration of glyphosate in biofilm tissues may be of particular concern for microalgal/photosynthetic organisms living in biofilms and consequently exposed to potentially harmful levels of glyphosate and AMPA, exceeding CCME guidelines, even if levels in the ambient water appear safe. Microalgae in biofilms are important contributors to shallow water food webs, oxygen and energy production via photosynthesis, and biogeochemical nutrient cycling (Battin et al., 2016). Microalgae and cyanobacteria exhibit variable and species-specific responses to glyphosate exposure (Choi et al., 2012; Forlani et al., 2008; Lozano et al., 2018; Smedbol et al., 2017; C. Wang et al., 2016). Cyanobacteria have been found to be tolerant to glyphosate at concentrations of ca. 0.03 mM (ca. 5 ppm) to >10 mM (ca. 1700 ppm) in some species, with the ability to metabolize and utilize glyphosate as a phosphorus source (Forlani et al., 2008; Huntscha et al., 2018; Ilikchyan et al., 2009), and low concentrations (0.1 mM, ca. 17 ppm) stimulating growth of some cyanobacteria (Berman et al., 2020; Drzyzga and Lipok, 2018). Conversely, cyanobacterial growth and photochemistry were found to be more sensitive to glyphosate compared to eukaryotic microalgae (Smedbol et al., 2017), with the half-maximal effective concentration (EC_50_) for growth ca. 400 μg·L^-1^ (ca. 0.4 ppm), while those for chlorophytes and a cryptophytes ranged from 400 – 1000 μg·L^-1^ (0.4 – 1 ppm). Concentrations producing negative responses were generally higher than the exposure levels within biofilm tissues observed here. Community responses can be further influenced by the availability of phosphorus when exposed to glyphosate (Berman et al., 2020; Carles et al., 2019; Huntscha et al., 2018; C. Wang et al., 2016), and the ability of different taxa to compete for and utilize this nutrient source may play a role.

Variable chlorophyll *a* fluorometry can provide an efficient, non-destructive method to measure algal responses to herbicide stress based on changes in Photosystem II and photochemical efficiency in phytoplankton (Choi et al., 2012; Smedbol et al., 2017) and periphyton (Chaumet et al., 2019; Dorigo and Leboulanger, 2001; Feckler et al., 2018; Tiam et al., 2015). Negative effects on biofilm photosynthetic efficiency have been observed at glyphosate concentrations on the order of 3 to >10 ppm (or mg·L^-1^), over varying exposure time periods (Bonnineau et al., 2012; Goldsborough and Brown, 1988; Iummato et al., 2017), yet we observed only a weak, non-significant trend of declining ΔF/F_m_’ with glyphosate exposure. Importantly, average post-exposure ΔF/F_m_’ was > 0.6 for all exposure concentrations, a quantum yield value typical of healthy algal cells (Campbell et al., 1998; Kolber et al., 1988). Glyphosate inhibits aromatic amino acid synthesis and does not directly target the photosynthetic apparatus. Hence, when additional stressors are absent and requirements for new cellular protein are minimal, it seems reasonable that brief exposures to 1-10 ppm glyphosate would have limited effects on the photosynthetic efficiency of biofilms. In contrast, maximum quantum yield (F_v_/F_m_) of freshwater phytoplankton was suppressed following glyphosate exposure (< 15 ppm (Choi et al., 2012); 0.5 – 1 ppm (Smedbol et al., 2017)), with responses following Michaelis-Menten saturation kinetics (Choi et al., 2012), but also showing considerable species-specific differences in sensitivity (Smedbol et al., 2017). Quantum yield alone may not be sensitive enough to assess herbicide stress in periphyton communities over short time scales (Dorigo and Leboulanger, 2001; Tiam et al., 2015), in particular for herbicides that do not directly affect Photosystem II (Feckler et al., 2018). Our results support this assessment, and we recommend parallel measurement of different responses over time to effectively capture herbicide effects on algal physiology and metabolism.

The range of response factors and species-specific variability suggests changes in photosynthetic community structure are likely following glyphosate exposure (Berman et al., 2020; Huntscha et al., 2018; Klátyik et al., 2017; Lozano et al., 2018; Pizarro et al., 2016; Smedbol et al., 2018). Changes to algal community composition have been observed at environmentally relevant concentrations of glyphosate (Magbanua et al., 2013), including increased abundance of chlorophytes (Klátyik et al., 2017) and cyanobacteria (Berman et al., 2020; Huntscha et al., 2018; Lozano et al., 2018; Pérez et al., 2007). We expect that chronic exposure to glyphosate may favour taxa with resistant forms of the target enzyme EPSPS or other tolerance mechanisms (Forlani et al., 2008; Huntscha et al., 2018), as well as those best able to utilize metabolically released phosphorus (Berman et al., 2020; Forlani et al., 2008; C. Wang et al., 2016). If changes in community composition negatively influence the health or abundance of species that are preferentially grazed by other organisms, indirect trophic effects may result. Increases in the abundance of cyanobacteria and/or planktonic algae may influence water quality and macrophyte abundance (Berman et al., 2020; Pizarro et al., 2016). This points to the need for future work examining the effects of chronic exposure on community structure along with biofilm functional characteristics, including autotrophy vs. heterotrophy (e.g. Feckler et al., 2018), pigment and lipid content. It would also be valuable to examine any changes in the relative abundance of type I and type II EPSPS pathways that would reveal a shift from sensitive to tolerant taxa and possibly an increase in the abundance of species with the C-P lyase necessary to harvest phosphorus from glyphosate.

### 4.1 Conclusions

The results presented reveal the ability of biofilms to metabolize glyphosate and retain and bioconcentrate glyphosate and its breakdown product AMPA. This demonstrates the importance of biofilms to improving water quality, facilitating contaminant removal from surface water and runoff – a valuable ecosystem function provided by wetlands and facilitated by biofilms. This is a potential explanation for the observed rapid dissipation of glyphosate from surface waters and the low levels detected even a short time after runoff events (Goldsborough and Beck, 1989; Goldsborough and Brown, 1993; Imfeld et al., 2013; Peruzzo et al., 2008). However, these same features present a potential negative impact, as biofilms also provide habitat and a food source for many invertebrates and juvenile aquatic organisms, including fish and amphibians (Battin et al., 2016). Bioconcentration of glyphosate and other pesticides in biofilms presents a contaminant delivery route to higher trophic levels that is not well understood (Lundqvist et al., 2012). The majority of ecotoxicological risk assessments examine physiological effects resulting from immersion, and we may be under-recognizing the potential ecological risk of contaminants, like glyphosate, that are bioconcentrating in biofilms and subsequently being consumed. Risk assessments for contaminants need to consider both the toxicity as well as the different exposure routes to organisms, and future ecotoxicity research should incorporate the effects of acute and chronic dietary exposure of glyphosate, as well as other contaminants, to aquatic biota.

## 5 Acknowledgements

Funding: This work was funded by research agreement NCA_20180507152929575 from the Ontario Ministry of Natural Resources and Forestry, National Science and Engineering Research Council of Canada Discovery Grant RGPIN-03846, and the Mitacs Career Connect program.

Field and laboratory assistance was provided by G. Howell, D. Anderson, H. Polowyk, S. Yuckin, H. Quinn-Austin, J. Pearson, L. Koiter, M. Tanguay and C. Robichaud. We thank __ anonymous reviewers for their feedback that improved this manuscript.

---

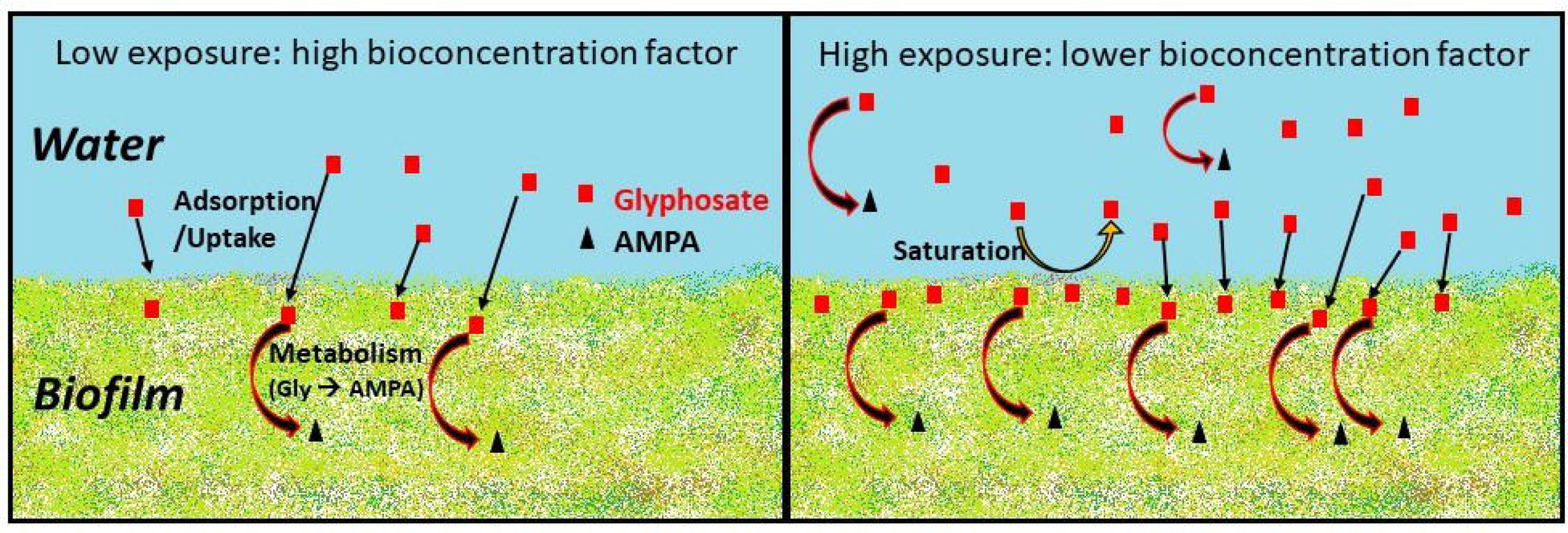

