## Supplementary materials for "Bioconcentration of glyphosate in wetland biofilms"

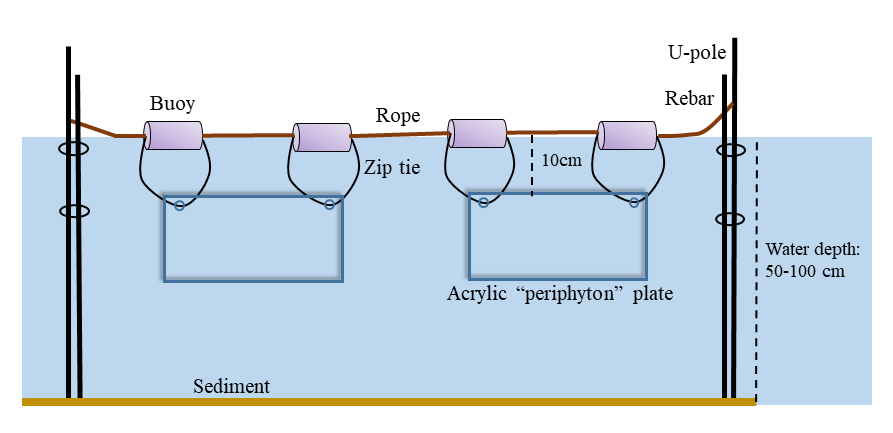

Figure S1. A schematic of the lay out of our biofilm sampling arrays. The schematic was reduced for display purposes, note that each array holds 4 or 5 acrylic plates.

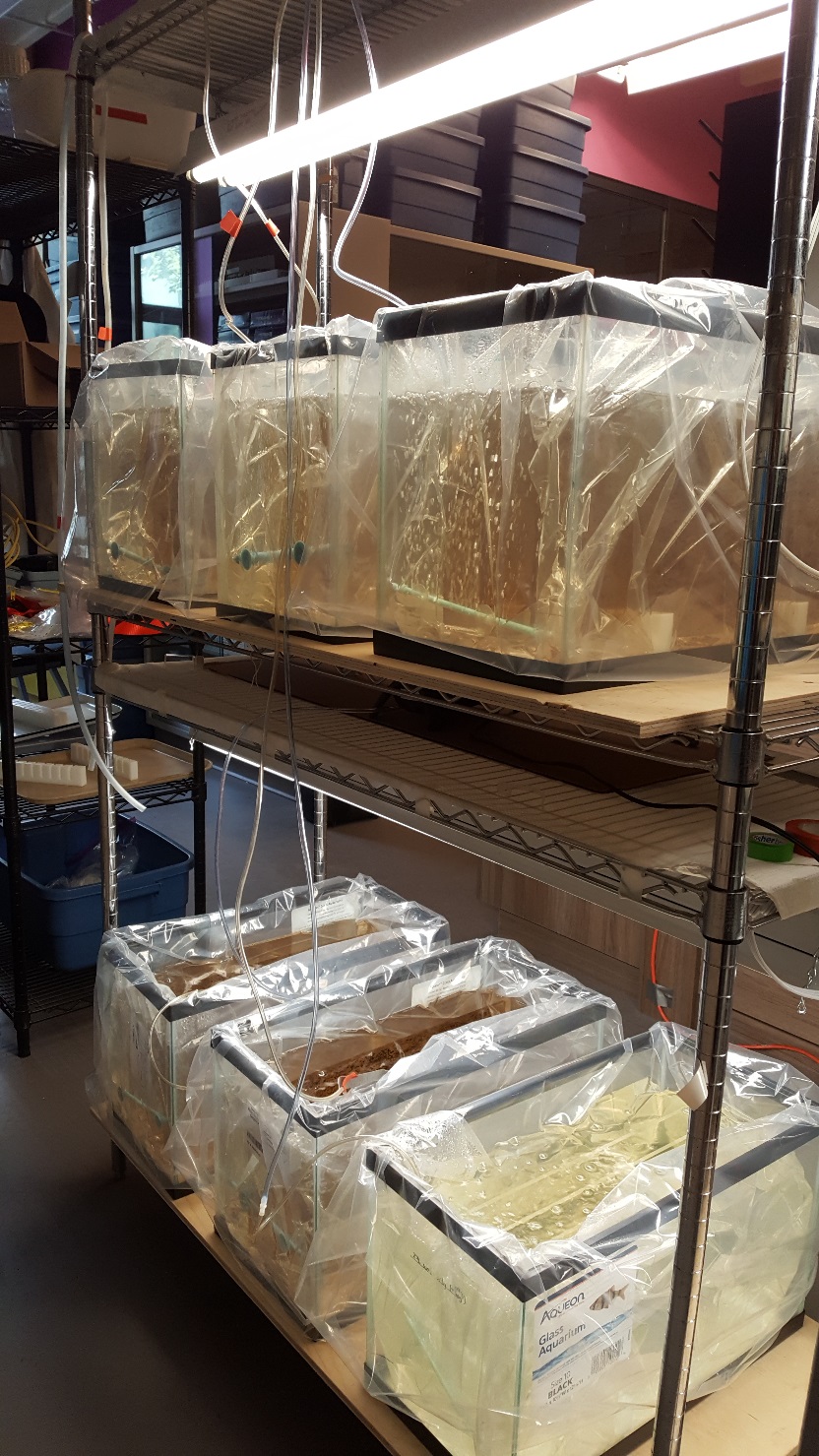

Figure S2. Laboratory Microcosm Set-up. Microcosms comprise glass aquaria lined with gusseted poly bags, aeration tubes, contain filtered site water, and are exposed to equal light exposure from cool white fluorescent lights. Five of the six microcosms contain colonized plates (darker colour) and one of the six contain un-colonized plates (clear).

Table S1. Analytic limits of detection (LOD) and quantification (LOQ) for glyphosate and amino-methyl phosphonic acid (AMPA) analyzed using liquid chromatography and mass spectrometry, as provided by the Agriculture and Food Laboratory (AFL), University of Guelph.

| Analyte | Sample matrix | Limit of Detection (ppm) | Limit of Quantification (ppm) |
| --- | --- | --- | --- |
| Glyphosate | water | 0.001 | 0.008 |
| AMPA | water | 0.002 | 0.008 |
| Glyphosate, AMPA | sediment | 0.005 | 0.02 |
| Glyphosate, AMPA | periphyton | 0.008 | 0.03 |

Table S2. Results of two-variable general linear models to assess if the relationship of bioconcentration factor (BCF) to ambient water concentration was different based on (a) application type (lab-dosed vs field-exposed) or (b) herbicide (AMPA vs. glyphosate).

|  |  | Estimate | Std. Error | t value | Pr(>\|t\|) |
| --- | --- | --- | --- | --- | --- |
| a | Intercept | 1.891 | 0.130 | 14.573 | 0.000 |
|  | Water concentration | -0.601 | 0.075 | -8.049 | 0.000 |
|  | Application type (factor) | -0.041 | 0.334 | -0.123 | 0.903 |
|  | Interaction | 0.250 | 0.190 | 1.314 | 0.205 |
| b | Intercept | 1.829 | 0.348 | 5.249 | 0.000 |
|  | Water concentration | -0.515 | 0.174 | -2.961 | 0.008 |
|  | Herbicide (factor) | 0.009 | 0.390 | 0.023 | 0.982 |
|  | Interaction | -0.045 | 0.205 | -0.217 | 0.831 |

There was no significant interaction or main effect from the application type or herbicide factors, therefore a single regression was used to estimate the relationship of BCF to ambient water concentration.

Table S3. Two-variable general linear model of the relationship between AMPA production (dependent) and glyphosate concentration (independent) with ‘application type’ (i.e. lab-dosed vs. field-exposed) as a factor.

|  | Estimate | Std. Error | t value | Pr(>\|t\|) |
| --- | --- | --- | --- | --- |
| Intercept | -0.705 | 0.877 | -0.804 | 0.429 |
| Glyphosate concentration | 0.282 | 0.034 | 8.232 | 1.39E-08 |
| Application type (factor) | -0.789 | 2.058 | -0.383 | 0.705 |
| Interaction | 0.068 | 0.048 | 1.43 | 0.165 |

There was no significant interaction between application type and glyphosate concentration on the amount of AMPA produced (p = 0.165) and no significant main effect of application type (p = 0.705), therefore application type was removed from the model. The intercept was forced through zero, as described in Methods, Statistical Analyses.

Table S4. Summary Parameters for Pre-exposure (time 0) and post-exposure (24 h post-dose) ΔF/F_m_^’^_­_ (light-adapted quantum yield of photochemistry) of biofilms in microcosms exposed to different glyphosate concentrations.

| **Glyphosate Conc. (ppm)** | **Pre-exposure** | | | | **Post-exposure** | | | |
| --- | --- | --- | --- | --- | --- | --- | --- | --- |
|  | **Mean** | **StDev** | **N** | **StError** | **Mean** | **StDev** | **N** | **StError** |
| 0 | 0.5598 | 0.1085 | 32 | 0.0192 | 0.6247 | 0.0332 | 36 | 0.0055 |
| 0.01 | 0.5969 | 0.0775 | 35 | 0.0131 | 0.6237 | 0.0291 | 34 | 0.0050 |
| 0.1 | 0.5575 | 0.1059 | 35 | 0.0179 | 0.6099 | 0.0214 | 34 | 0.0037 |
| 1 | 0.5931 | 0.0636 | 35 | 0.0107 | 0.6146 | 0.0490 | 35 | 0.0083 |
| 10 | 0.6040 | 0.0510 | 34 | 0.0088 | 0.6105 | 0.0338 | 34 | 0.0058 |

Normalized Difference in ΔF/Fm’­

| Glyphosate Concentration (ppm) | Normalized Post Y |
| --- | --- |
| 0 | 0.115949 |
| 0.01 | 0.0450328 |
| 0.1 | 0.0940929 |
| 1 | 0.0361272 |
| 10 | 0.0107621 |

| Glyphosate Concentration (ppm) | Regn Estimates |
| --- | --- |
| 0 | 0.075455 |
| 0.01 | 0.0753872 |
| 0.1 | 0.0747772 |
| 1 | 0.068677 |
| 10 | 0.007675 |
| 11 | 0.000897 |

Figure 3. Difference in average post-exposure ΔF/F_m_^’^_­_ (light-adapted quantum yield) compared to average pre-exposure values, normalized to pre-exposure values, at different glyphosate concentrations. Linear regression analysis (dashed line) indicated a non-significant negative relationship of change in ΔF/F_m_^’^_­_ with glyphosate concentration (y = -0.0068*x* + 0.0755, F_1,3_ = 2.628, p = 0.2034, adjusted r^2^ = 0.2893, RSE = 0.03653).
